# Bioinformatic analysis reveals the molecular mechanisms of Trefoil factor 3 involved in asthma pathogenesis

**DOI:** 10.1101/2025.01.20.633844

**Authors:** Fang He, Shu Wang, Pengfei Huang, Xiaoyun Fan

## Abstract

**Objective:** Investigate the expression, regulatory network and function of TFF3 in asthma from a multi-omics perspective through bioinformatics and cellular experiments to provide clues for understanding its molecular mechanisms in asthma.

**Methods:** Downloaded two asthma-related datasets (GSE67472 and GSE147878) from GEO. Conducted differential expression analysis and WGCNA to find common genes related to asthma phenotype and TFF3 expression. Constructed a PPI network to identify key genes interacting with TFF3. Performed GO and pathway enrichment analysis using DAVID. Analyzed the relationship between TFF3 and immune cell infiltration with CIBERSORTx. Used molecular docking to validate interactions. In vitro, induced 16HBE cells with HDM to establish asthma model and detected inflammatory indicators and TFF3 by RT-qPCR and western blotting.

**Results:** TFF3 expression was significantly increased in asthma patients in both datasets. ROC analysis showed good diagnostic specificity and sensitivity. 57 co-expressed genes related to asthma and TFF3 were screened, with 11 genes directly interacting with TFF3 and upregulated. Functional enrichment analysis indicated involvement in asthma-related processes. Immune infiltration analysis showed increased M2 macrophages and mast cells, decreased M1 macrophages, and positive correlation with TFF3. Molecular docking confirmed stable binding. In vitro experiments showed increased inflammation index and TFF3 expression after HDM intervention.

**Conclusion:** The study used bioinformatics and cellular experiments to show TFF3’s key roles in asthma pathogenesis in aspects like gene expression, protein interaction, and immune microenvironment. TFF3 may regulate genes like AGR2 and AKT1, affect immune cells, and be involved in processes like cell migration and immune response. It provides clues for exploring molecular mechanisms and may offer new directions for asthma diagnosis and treatment.

## 1 Introduction

Asthma is a common chronic respiratory disease affecting approximately 334 million people worldwide and seriously impacting patients’ physical and mental health and quality of life [1]. he main pathological features of asthma include chronic airway inflammation, airway hyperresponsiveness, and airway remodeling, involving complex pathogenic mechanisms and multiple cells and inflammatory mediators [2]. Although several treatment options exist, such as inhaled corticosteroids and β2 receptor agonists, a considerable proportion of patients still respond poorly to current therapies [2]. Therefore, it is of great significance to further explore the pathogenesis of asthma and identify new therapeutic targets and strategies.TFF3 (Trefoil factor 3) is a secreted protein highly expressed in respiratory epithelial cells [3, 4]. Multiple studies in recent years have demonstrated its close relationship with the occurrence and development of asthma. For instance, researchers have found significantly elevated TFF3 levels in the serum and sputum of asthma patients compared to healthy individuals and showed upregulated TFF3 expression in asthmatic bronchial epithelial cells [5, 6]. TFF3 can be involved in asthma’s pathological process by regulating mucin secretion and promoting epithelial cell differentiation and proliferation [3]. Inhibiting TFF3 expression with budesonide can improve airway inflammation and mucus secretion in acute allergic inflammation mice [7]. These studies emphasize the important role of TFF3 in asthma but its specific mechanism and regulatory mechanisms remain unclear and need further elucidation.

Bioinformatics analysis provides a powerful tool to investigate gene functions in diseases. By integrating high-throughput data from different sources, bioinformatics analysis can be used to screen key disease-related genes, construct molecular regulatory networks, and thereby reveal disease molecular mechanisms [8]. In recent years, multiple studies have used bioinformatics methods to identify some key genes and their regulatory mechanisms related to asthma, such as CH25H, CAMKK2, etc [9, 10]. However, there is still a lack of systematic bioinformatic analysis on the mechanism of action of TFF3 in asthma.

This study aims to systematically explore TFF3’s expression pattern, regulatory network and function in asthma using bioinformatics methods based on public asthma transcriptome data. The goal is to provide clues for understanding TFF3’s molecular mechanisms in asthma pathogenesis and new ideas for asthma diagnosis and treatment.

## 2 Methods

### 2.1 Data download and preprocessing

Two asthma-related gene expression profile datasets, GSE67472 and GSE147878, were downloaded from the Gene Expression Omnibus (GEO) database (https://www.ncbi.nlm.nih.gov/geo/) at NCBI. The GSE67472 dataset based on the GPL16311 platform included airway epithelial tissue samples obtained by bronchoscopy from 43 healthy controls (Normal control, NC) and 62 asthma patients (Asthma, AA). The GSE147878 dataset based on the GPL10558 platform included airway epithelial tissue samples obtained by bronchoscopy from 13 healthy controls (NC) and 60 asthma patients (AA).

First, gene name mapping was done on the probes as per the platform annotation file. Duplicate probe expression values were removed. When multiple probes corresponded to one gene, the average value was taken as that gene’s expression level. Then, the limma package was employed for background correction and data normalization. The quantile normalization method was used for inter-group normalization to get rid of systematic bias from technical factors. The data was converted to log2 values for subsequent analysis.

### 2.2 TFF3 expression analysis and ROC analysis

Based on the GSE67472 and GSE147878 datasets respectively, the Wilcoxon rank-sum test was used to compare the expression differences of TFF3 in healthy airway epithelial tissues and asthma patient airway tissue samples, and box plots were drawn. In addition, receiver operating characteristic (ROC) curve analysis was used to evaluate the diagnostic value of TFF3 for asthma, and the area under the curve (AUC) value was used to measure the diagnostic efficacy.

### 2.3 Screening of genes related to TFF3 expression in asthma

Based on the GSE67472 and GSE147878 datasets respectively, genes related to TNFSF11 expression in osteoarthritis were identified through differential expression analysis and weighted gene co-expression network analysis (WGCNA). Specifically, differentially expressed genes in airway epithelial tissue samples from the AA group and NC group were screened using thresholds of “|Fold change|=1.2, *P*<0.05”. In WGCNA, we first constructed an adjacency matrix to describe the association strength between nodes. Then, the optimal soft threshold β was selected to transform the adjacency matrix into a topological overlap matrix, so that the constructed network conforms to a power-law distribution and is closer to a real biological network state. Second, the blockwiseModules function was used to construct a scale-free network, and then module division was performed using average linkage hierarchical clustering and dynamic tree cutting algorithms to determine gene co-expression modules. Then, the eigengenes (MEs) of each gene module were calculated and the correlation between each module and clinical grouping (AA group) and TFF3 expression was analyzed to screen gene modules significantly correlated with both clinical grouping (AA group) and TFF3 expression. Then, by calculating the gene significance (GS) of each gene in these gene modules related to clinical grouping (AA group) and TFF3 expression, hub genes of the co-expression network were screened using a threshold of GS>0.5. Finally, by comparing the differentially expressed genes and co-expression network hub genes in the GSE67472 and GSE147878 datasets, the common genes were considered to be genes related to TFF3 expression in asthma.

### 2.4 PPI network analysis and functional enrichment analysis

To explore the co-expressed genes with potential mutual regulatory effects with TFF3 in asthma, we used the STRING database (https://cn.string-db.org/) to perform protein-protein interaction (PPI) analysis on the “genes related to TFF3 expression in asthma” identified in 2.3. Then, Cytoscape software was used for visual analysis and topological structure analysis of the PPI network to identify genes that directly and indirectly interact with TFF3 in asthma. The non-parametric Wilcoxon rank-sum test was then used to compare the expression differences of genes directly interacting with TFF3 in the PPI network between the AA group and NC group. Pearson correlation analysis was used to study the expression correlation between these genes and with TNFSF11.

To further explore the biological functions involved in TFF3’s participation in the occurrence and development of asthma, we used the DAVID database (https://david.ncifcrf.gov) to perform Gene Ontology (GO) and pathway enrichment analysis on all genes in the PPI network, and functional enrichment analysis on TFF3 and its directly related genes in the PPI network. Fisher’s exact test was used for statistical testing, with *P*<0.05 indicating significant enrichment.

In addition, we used the “gsea” package in R software to perform Gene Set Enrichment Analysis (GSEA) to further compare the expression differences of signaling pathways involved in TFF3 and its directly related genes between the AA and NC groups in the two datasets, in order to verify the accuracy of the functional enrichment analysis. The normalized enrichment score (NES) and P-value were used to evaluate the expression changes of related pathways, with |NES|>1 and *P*<0.05 indicating significant expression differences in the signaling pathway.

### 2.5 Molecular docking analysis

Molecular docking technology was further used to verify the interaction between TFF3 and the proteins encoded by its directly related genes in the PPI network. Specifically, the crystal structures of TFF3 and the proteins encoded by its directly related genes were first downloaded from the RCSB Protein Data Bank (https://www.rcsb.org/). The HDOCK protein-protein docking server (http://hdock.phys.hust.edu.cn/) was used for molecular docking analysis. After docking, the binding energy was used to evaluate the binding ability of the complex, with higher negative values of binding energy indicating stronger binding ability. The Confidence Score was used to evaluate the docking confidence and the Ligand RMSD (L-RMSD) value was used to evaluate the conformational change, with larger Confidence Scores indicating higher docking confidence and smaller L-RMSD indicating smaller conformational changes and more stable binding. Finally, the PyMOL molecular graphics system software (version 2.3) was used to visually analyze the screened complexes and observe the interaction patterns such as hydrogen bonds, hydrophobic interactions, and van der Waals forces to analyze the binding patterns between TFF3 and its direct target proteins.

### 2.6 Immune infiltration analysis

we used the CIBERSORTx (https://cibersortx.stanford.edu/) database to analyze the types and relative abundance of immune cells in airway epithelial tissues of asthma patients (AA group) and healthy airway epithelial tissue samples (NC group) in the GSE67472 and GSE147878 datasets. *P*<0.05 was the criterion for significant infiltration differences. Pearson correlation analysis was used to analyze the correlation between the infiltration level of each immune cell type and TFF3 expression.

### 2.7 Cell culture and cell processing

16HBE (human bronchial epithelial cell) cell line and HAM (human alveolar macrophage) cell line were obtained from Qingqi Biotechnology Development Co Ltd (Shanghai). All cell lines were characterized using short tandem duplicate assays. Cells were cultured in DMEM complete medium at 37°C with 5% CO2. The cells were divided into control group and HDM group. The HDM group was stimulated in medium with concentrations of 200, 400, and 800 U/mL HDM and cultured for 24 h. Cells were subsequently collected for assay.

### 2.8 Western blot

Total protein was obtained using the Total Protein Extraction Kit (ProteinTech) according to the manufacturer’s instructions. Proteins were separated by electrophoresis on 8-15% sodium dodecyl sulfate-polyacrylamide gels and transferred to PVDF membranes. After being closed with 5% skim milk powder for 1h at room temperature, the membranes were incubated overnight at 4°C with the appropriate antibodies, including: TFF3 (abcam), AKT (CST), phosphorylated AKT (p-AKT) (CST), PI3K (CST), phosphorylated PI3K (p-PI3K) (CST) and GAPDH (ProteinTech). After incubation of PVDF membranes with appropriate secondary antibodies, visualization was performed using the Tanon 5200 system and ECL detection reagents and quantified using ImageJ software.

### 2.9 Real Time-qPCR

Total RNA from cells was extracted using TRIzol. cDNA was synthesized using a reverse transcription (Thermo Fisher Scientific) kit according to the manufacturer’s instructions. Quantitative PCR was performed using a universal SYBR green rapid qPCR mixture (Yeasen Biotechnology) in a fluorescence quantitative real-time PCR instrument (Bio Rad). GAPDH was used as the reference gene. Primer sequences were shown in supplementary Table 1.

## 3 Results

### 3.1 TFF3 expression analysis and ROC analysis

Expression analysis results showed that compared with the NC group, TFF3 expression was significantly upregulated in the AA group in both the GSE67472 and GSE147878 datasets (*P*<0.05) (Figure 1A, 1B). ROC analysis results showed that in the GSE67472 and GSE147878 datasets, the area under the curve (AUC) of TFF3 was 0.791 and 0.842, respectively (Figure 1C, 1D). These results indicate that TFF3 has good diagnostic value for asthma and plays an important role in the development of asthma, which is consistent with previous reports.

**Figure 1.**
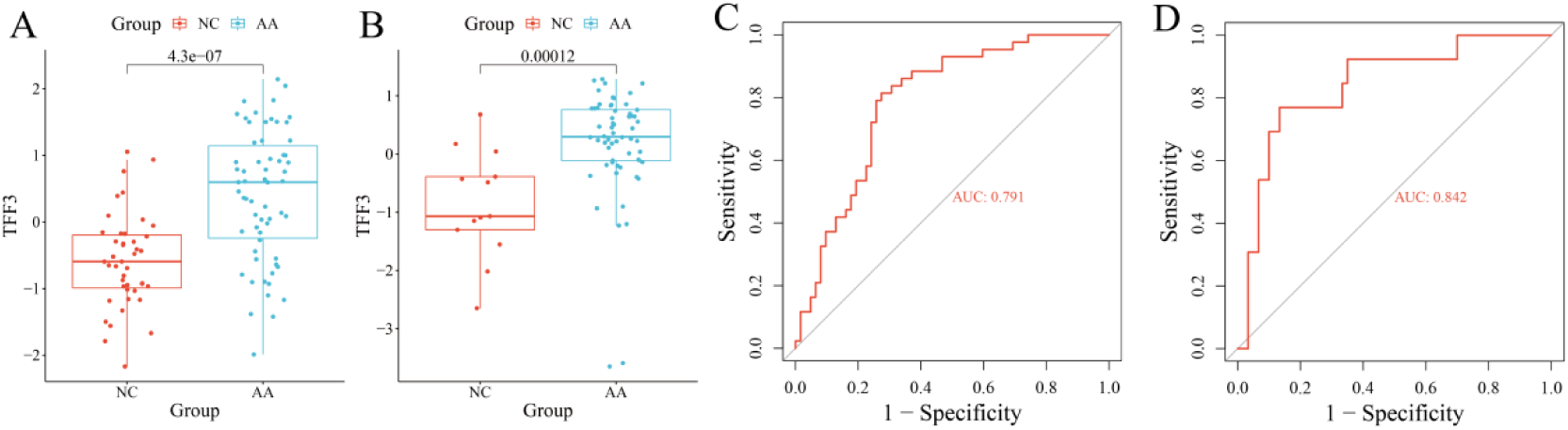
TFF3 expression analysis and ROC analysis in asthma. (A, B) Box plots showing the expression levels of TFF3 in airway epithelial tissues of the AA and NC groups in the GSE67472 and GSE147878 datasets. (C, D) ROC curves evaluating the diagnostic value of TFF3 expression for asthma in the GSE67472 and GSE147878 datasets.

### 3.2 Identification of genes related to TFF3 expression in asthma

First, genes related to TFF3 expression were screened based on the GSE67472 dataset. Differential expression analysis results showed that 973 differentially expressed genes between the AA and NC groups were screened (|Fold Change|>1.2, *P*<0.05), including 633 genes significantly upregulated in the AA group and 340 genes significantly downregulated in the AA group (Figure 2A, 2B). WGCNA analysis results showed that by setting the optimal soft threshold β to 8, a total of 20 gene co-expression modules were generated (Figure 2C). These modules have good independence, and the similarity connectivity distances between them are all greater than 0.25 (Figure 2D). Among them, 7 gene modules (darkgrey, lightyellow, blue, greenyellow, cyan, orange, turquoise) were significantly correlated with both clinical grouping (AA group) and TFF3 expression (Figure 2E). Using GS>0.5 as the threshold, a total of 2899 hub genes were identified from these significant gene modules. By comparing the differentially expressed genes and hub genes, a total of 745 genes related to asthma phenotype and TFF3 expression were obtained (Figure 2F).

**Figure 2.**
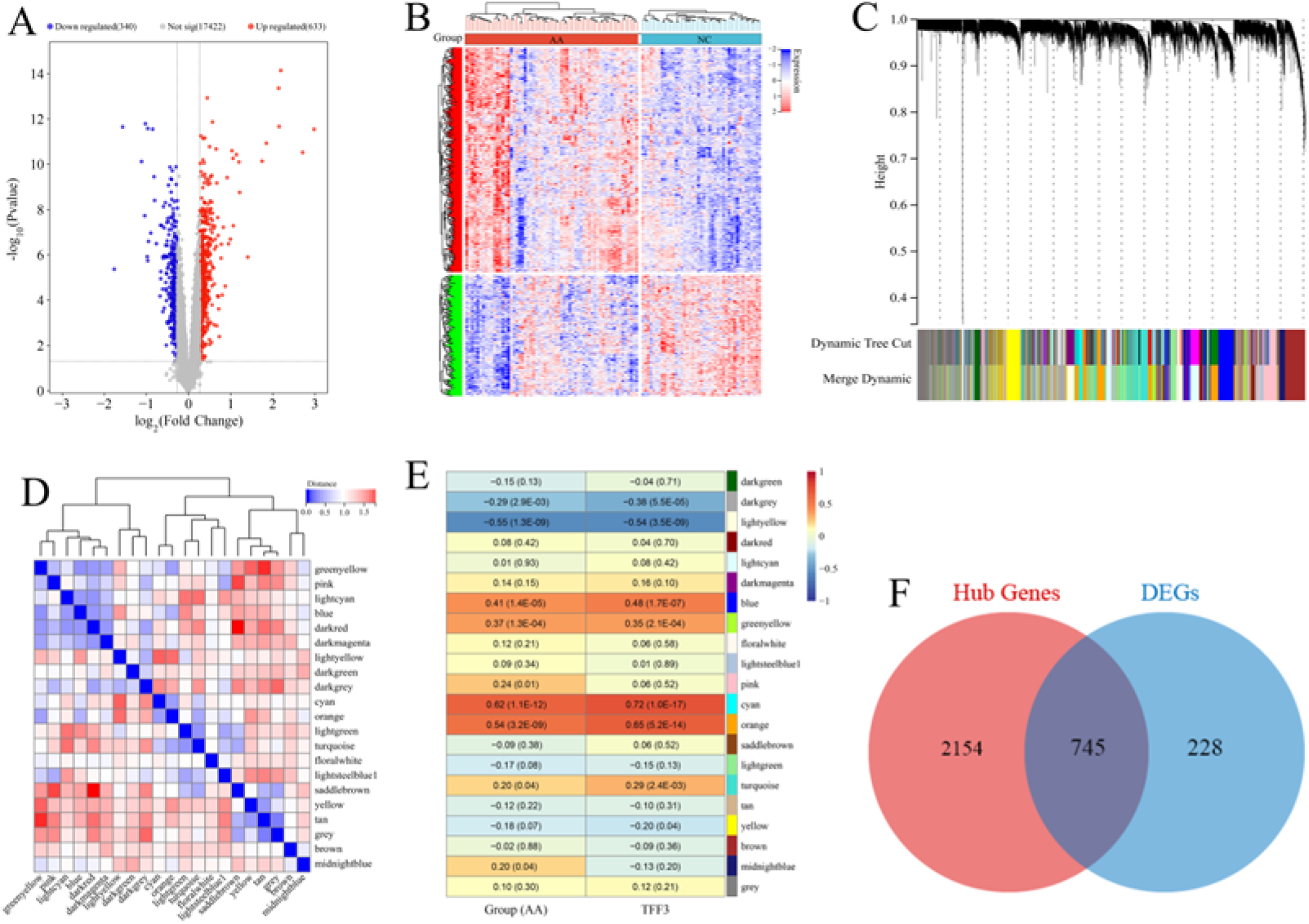
Screening of genes related to TFF3 expression in asthma based on the GSE67472 dataset. (A) Volcano plot of differentially expressed genes between the AA and NC groups. Red indicates significantly upregulated genes in the AA group, while green indicates significantly downregulated genes (|Fold Change|>1.2, P<0.05). (B) Heatmap of differentially expressed genes between the AA and NC groups. (C) Gene dendrogram obtained by average linkage hierarchical clustering. The color row underneath the dendrogram shows the module assignment determined by the Dynamic Tree Cut. (D) Heatmap of the adjacencies in the gene network. (E) Heatmap of the correlation between module eigengenes and clinical traits (asthma status and TFF3 expression). (F) Venn diagram showing the overlap of differentially expressed genes and hub genes in WGCNA.

Then, genes related to TFF3 expression were screened based on the GSE147878 dataset. Differential expression analysis results showed that 755 differentially expressed genes between the AA and NC groups were screened (318 genes significantly upregulated in the AA group, 437 genes significantly downregulated in the AA group) (Figure 3A, 3B). In WGCNA analysis, the optimal soft threshold β was set to 5, and a total of 22 independent gene co-expression modules were generated (Figure 3C, 3D). 6 gene modules (violet, darkmagenta, cyan, salmon, darkgrey, lightcyan1) were significantly correlated with both clinical grouping (AA group) and TFF3 expression (Figure 3E). Using GS>0.5 as the threshold, a total of 2640 hub genes were identified from these significant gene modules. By comparing the differentially expressed genes and hub genes, a total of 311 genes related to asthma phenotype and TFF3 expression were obtained (Figure 3F).

**Figure 3.**
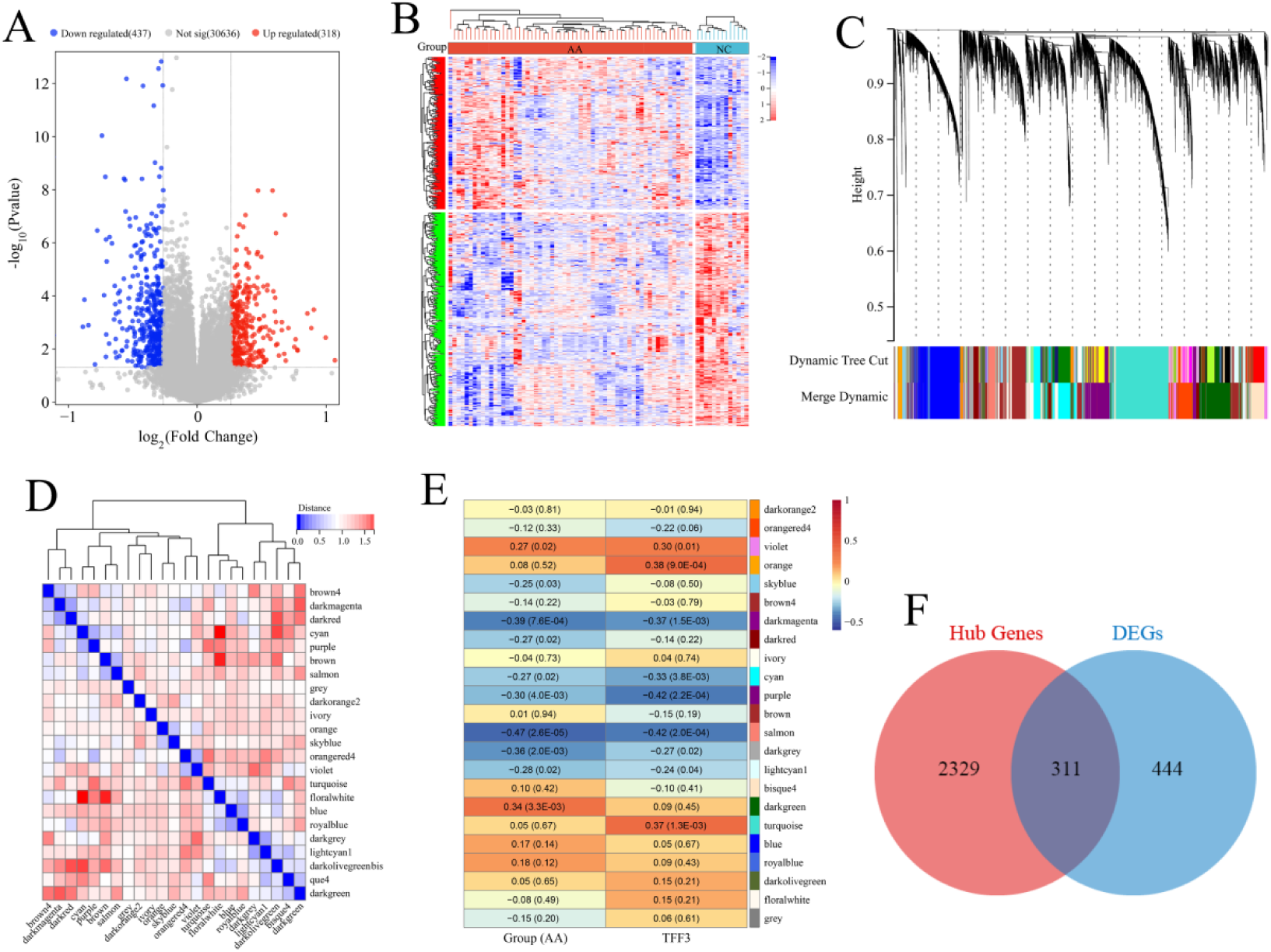
Screening of genes related to TFF3 expression in asthma based on the GSE147878 dataset. (A) Volcano plot of differentially expressed genes between the AA and NC groups. (B) Heatmap of differentially expressed genes between the AA and NC groups. (C) Gene dendrogram obtained by average linkage hierarchical clustering. (D) Heatmap of the adjacencies in the gene network. (E) Heatmap of the correlation between module eigengenes and clinical traits (asthma status and TFF3 expression). (F) Venn diagram showing the overlap of differentially expressed genes and hub genes in WGCNA.

### 3.3 PPI network analysis

By comparing the GSE67472 and GSE147878 datasets, 57 common genes related to asthma phenotype and TFF3 expression were finally screened (Figure 4). Protein-protein interaction analysis was performed on TFF3 and its 57 co-expressed genes, and the results are shown in Figure 4A. The PPI network (Figure 4B) contains a total of 54 nodes and 173 edges. Among them, 11 nodes (AGR2, AKT1, ATP10B, CLDN10, CPA3, CXCL14, DEFB1, MS4A2, S100P, SCGB2A1, SPDEF) have direct interaction with TFF3, and 42 nodes have indirect interaction with TFF3, indicating that TFF3 may directly or indirectly regulate these genes to participate in the development of asthma. These genes that directly interact with TFF3 may be key regulatory factors for TFF3 to participate in the development of asthma. We further analyzed the expression differences of these TFF3 direct interaction genes in the AA and NC groups and analyzed the expression correlation between these genes and TFF3. The results showed that compared with the NC group, these genes were all significantly upregulated in the AA group (*P*<0.05) (Figure 4C, 4D); all genes had significant positive correlations with TFF3 (R>0, *P*<0.05) (Figure 4E, 4F).

**Figure 4.**
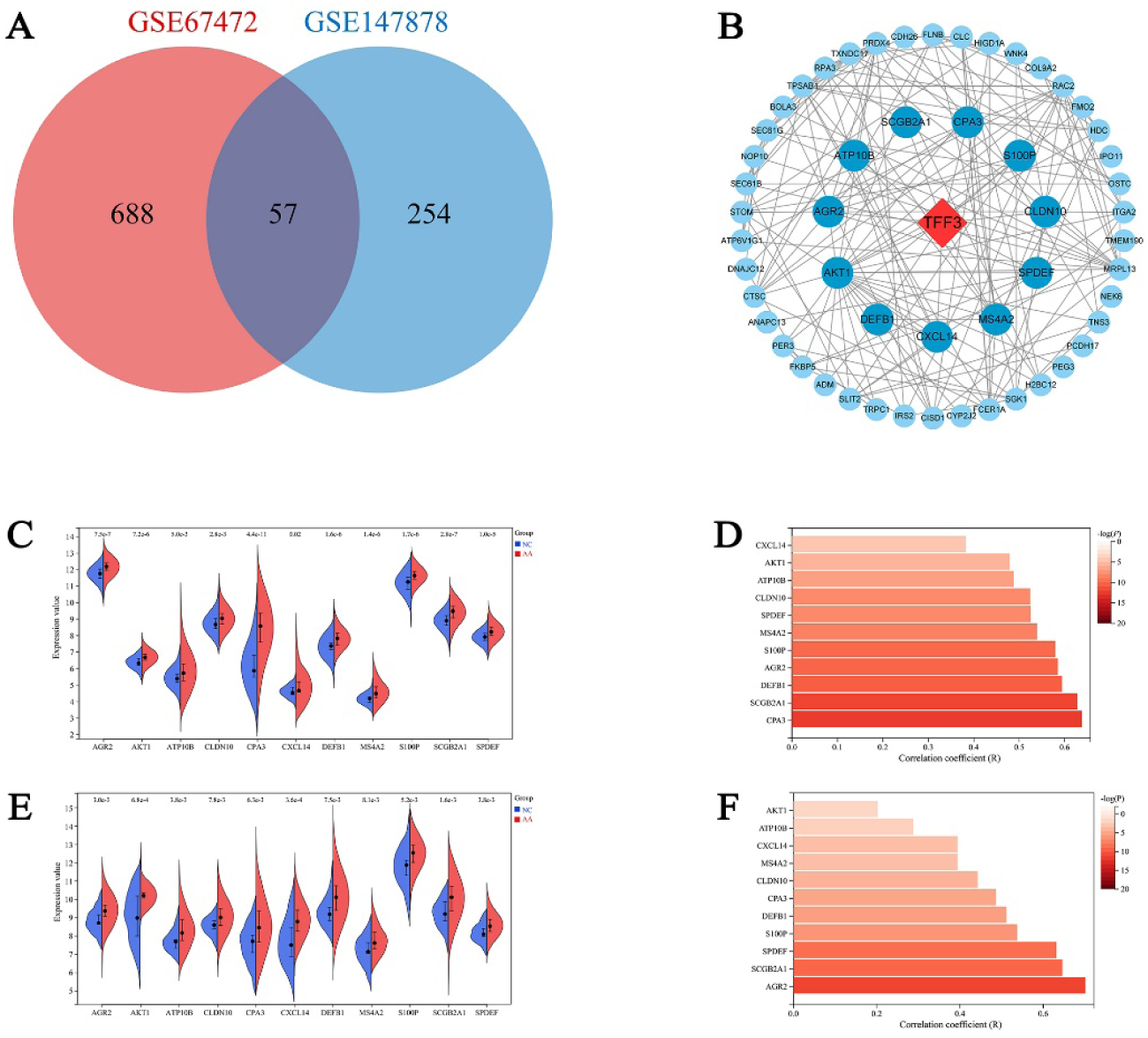
Venn diagram showing the overlap of genes related to asthma phenotype and TFF3 expression in the GSE67472 and GSE147878 datasets. (A) Venn diagram showing the overlap of genes related to asthma phenotype and TFF3 expression in the GSE67472 and GSE147878 datasets. (B) PPI network of TFF3 and its co-expressed genes. AGR2, AKT1, ATP10B, CLDN10, CPA3, CXCL14, DEFB1, MS4A2, S100P, SCGB2A1, SPDEF are targets of direct interaction with TFF3, while other targets are targets of indirect interaction with TFF3. (C, D) Expression levels of TFF3 direct interaction genes in the AA and NC groups in the GSE67472 and GSE147878 datasets. (E, F) Pearson correlation analysis of the expression levels between TFF3 and its direct interaction genes in the GSE67472 and GSE147878 datasets.

Then, we explored the biological functions involved in TFF3’s participation in the development of asthma. The results showed that the GO functional annotations of TFF3 and its interacting genes included 12 biological processes such as Regulation of cell migration, Lung goblet cell differentiation, Negative regulation of plasma membrane long-chain fatty acid transport, Regulation of mitotic cell cycle, and Negative regulation of monocyte chemotaxis; 5 cellular components such as Extracellular space, Extracellular exosome, Extracellular region, Endoplasmic reticulum membrane, and Focal adhesion; and 6 molecular functions such as Protein binding, Identical protein binding, IgE binding, Chloride ion binding, and Protein homodimerization activity (Figure 5A). The signaling pathways involved in these genes include 8 KEGG signaling pathways such as Fc epsilon RI signaling pathway, Sphingolipid signaling pathway, Asthma, Phospholipase D signaling pathway, and Focal adhesion; 7 REACTOME signaling pathways such as Ion channel transport, Immune System, Stimuli-sensing channels, Innate Immune System; and 8 WIKIPATHWAYS signaling pathways such as Insulin signaling, Focal adhesion, MECP2 and associated Rett syndrome, Integrin mediated cell adhesion, and Leptin insulin signaling overlap (Figure 5B). In addition, we also performed pathway enrichment analysis on TFF3 and its direct interaction genes. The results showed that these genes are involved in 4 KEGG signaling pathways (Fc epsilon RI signaling pathway, Sphingolipid signaling pathway, Phospholipase D signaling pathway, Asthma), 3 REACTOME pathways (Esr mediated signaling, Signaling by nuclear receptors, Innate immune system) and 2 WIKIPATHWAYS pathways (Epithelial to mesenchymal transition in colorectal cancer, Chemokine signaling pathway) (Figure 5C, 5D). These results suggest that TFF3 may participate in the above biological functions and signaling pathways by regulating other genes, thereby promoting the development of asthma.

**Figure 5.**
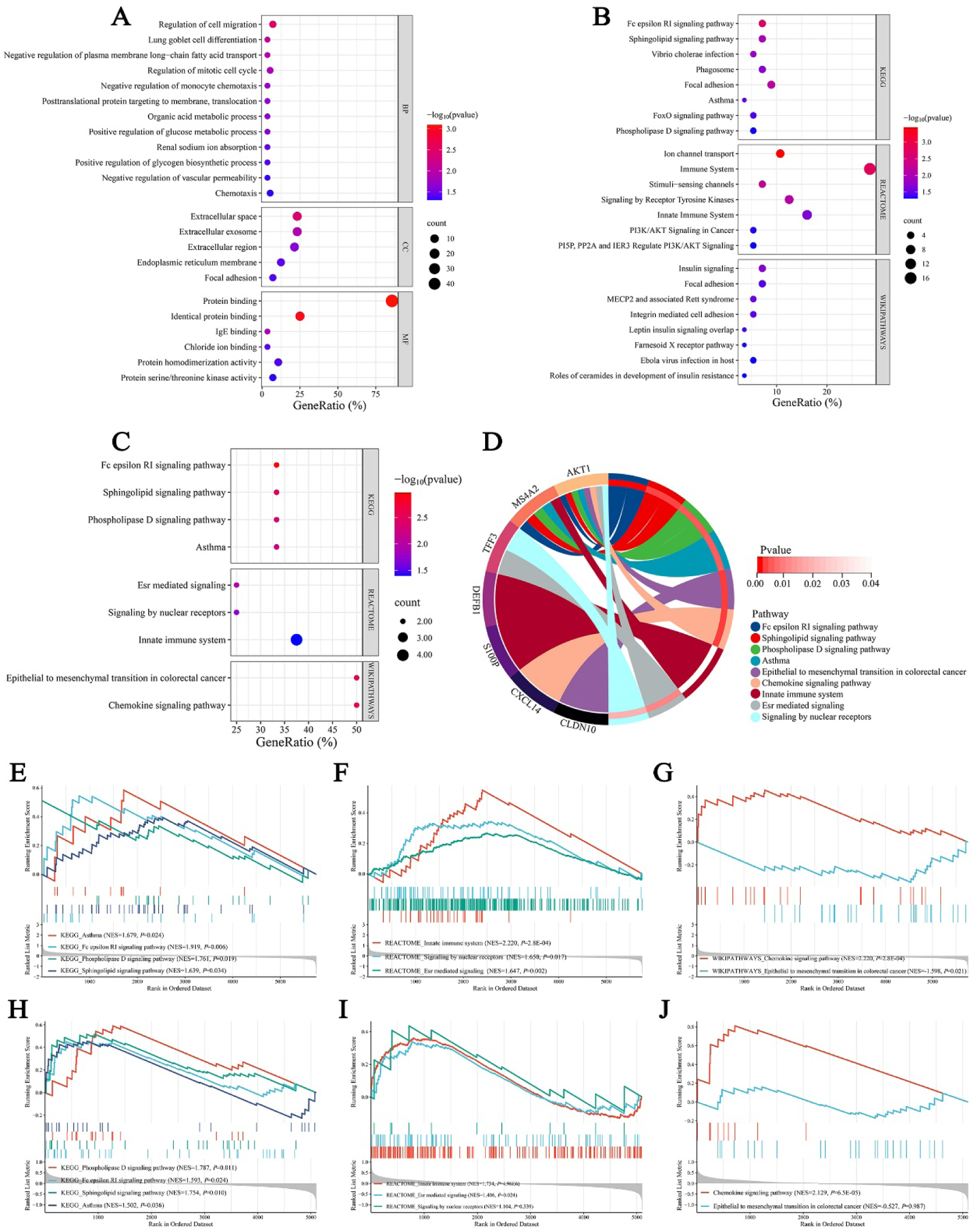
Functional enrichment analysis of TFF3 and its interacting genes. (A) GO annotation of TFF3 and its interacting genes. (B) KEGG, REACTOME and WIKIPATHWAYS pathway enrichment of TFF3 and its interacting genes. (C) KEGG, REACTOME and WIKIPATHWAYS pathway enrichment of TFF3 and its direct interaction genes. (D) Chord diagram showing the correlation between enriched pathways and related genes of TFF3 direct interaction genes. GSEA results comparing signaling pathway differences between the AA and NC groups in the GSE67472 and GSE147878 datasets. (E-G) GSEA results based on KEGG, REACTOME and WIKIPATHWAYS gene sets in the GSE67472 dataset, respectively. (H-J) GSEA results based on KEGG, REACTOME and WIKIPATHWAYS gene sets in the GSE147878 dataset, respectively.

To verify the accuracy of functional enrichment analysis results, the GSEA method was used to compare the expression differences of signaling pathways related to TFF3 and its directly related genes in the AA (asthma group) and NC (normal control group) of two datasets. The results showed that in the GSE67472 and GSE147878 datasets, several pathways including Fc epsilon RI signaling pathway, Sphingolipid signaling pathway, etc. were significantly enriched in the AA group compared to the NC group (Figure 5E-5J). These results are consistent with functional enrichment analysis, further confirming that TFF3 may be involved in asthma’s occurrence and development by regulating these signaling pathways.

### 3.4 Molecular docking analysis

Molecular docking technology was further used to verify the interaction between the proteins encoded by genes directly related to TFF3 in the PPI network and the TFF3 protein. The results showed that the proteins encoded by these genes directly related to TFF3 in the PPI network (AGR2, AKT1, ATP10B, CLDN10, CPA3, CXCL14, DEFB1, MS4A2, S100P, SCGB2A1, SPDEF) had good binding ability with the TNFSF11 protein (Confidence Score>0.8, binding energy<-200 kcal/mol), indicating a strong binding interaction (Table 2,in supplementary). In addition, we performed a visual analysis of the “ligand-receptor” complexes to analyze the interaction patterns between TFF3 and its ligands. The results demonstrated that the TFF3 protein could interact with AGR2, AKT1, ATP10B, CLDN10, CPA3, CXCL14, DEFB1, MS4A2, S100P, SCGB2A1, and SPDEF proteins by forming stable hydrogen bonds and salt bridges (Figure 6).

**Figure 6.**
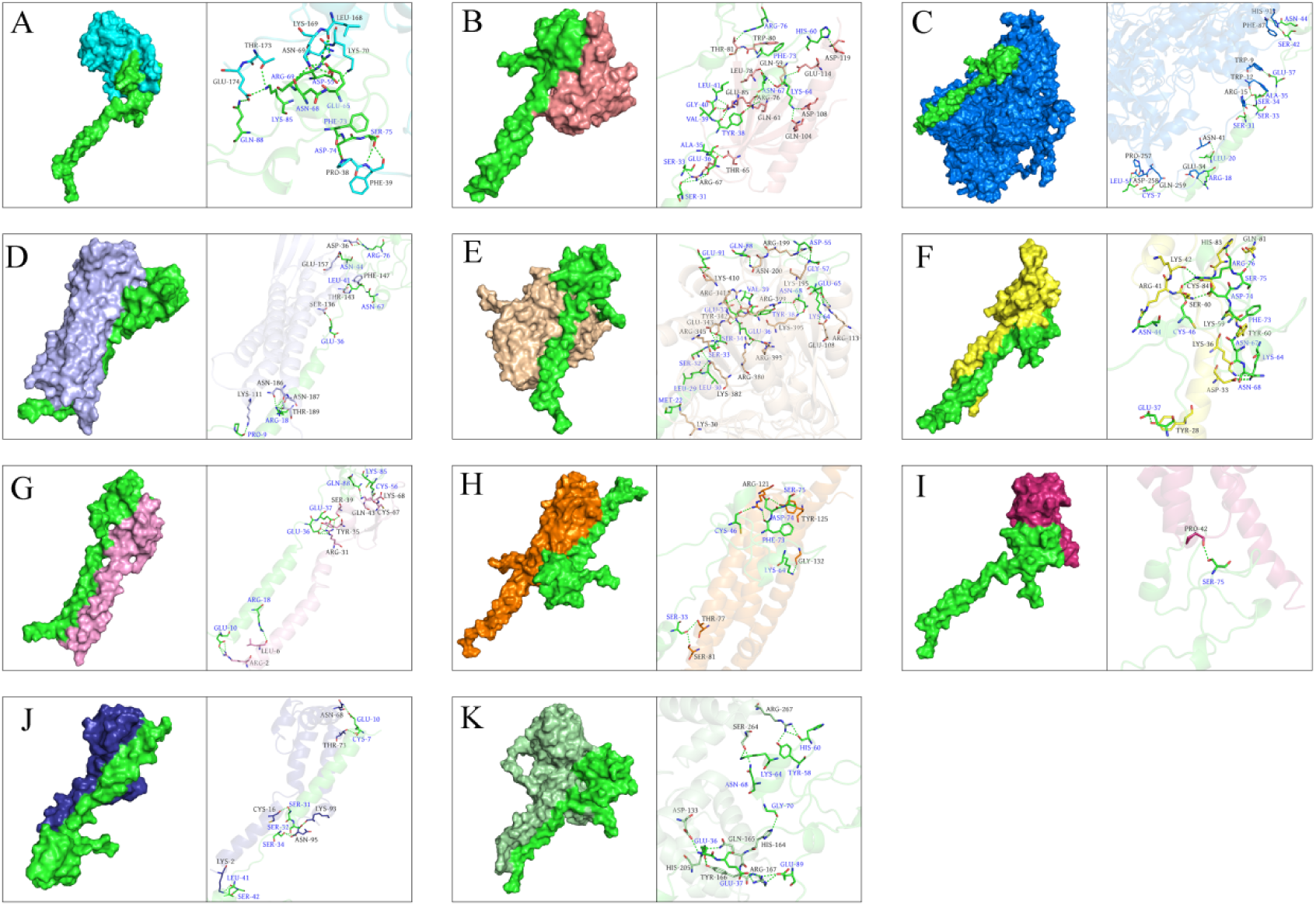
Molecular docking analysis of TFF3 protein with proteins encoded by its direct interaction genes. (A-K) Interaction patterns of TFF3 protein with AGR2, AKT1, ATP10B, CLDN10, CPA3, CXCL14, DEFB1, MS4A2, S100P, SCGB2A1, and SPDEF proteins.

### 3.5 Immune infiltration analysis

Immune cell infiltration plays a crucial role in the occurrence and development of asthma. In this study, we further explored the correlation between TFF3 and immune cell infiltration in asthma. The types and quantities of immune cell infiltration in each sample of the GSE67472 and GSE147878 datasets are presented in Figure 7A and Figure 7B. In both datasets, compared with the NC group, the infiltration levels of Macrophages M2 and Mast cells activated were significantly increased, while the infiltration of Macrophages M1 was significantly decreased in the AA group (Figure 7C, 7D). The expression of TFF3 was significantly negatively correlated with the infiltration levels of Mast cells resting and Macrophages M0, and significantly positively correlated with the infiltration levels of Macrophages M2 and Mast cells activated (Figure 7E, 7F). These results suggest that TFF3 may promote the occurrence and development of asthma by enhancing the immune infiltration of Macrophages M2 and Mast cells activated and reducing the number of Macrophages M0.

**Figure 7.**
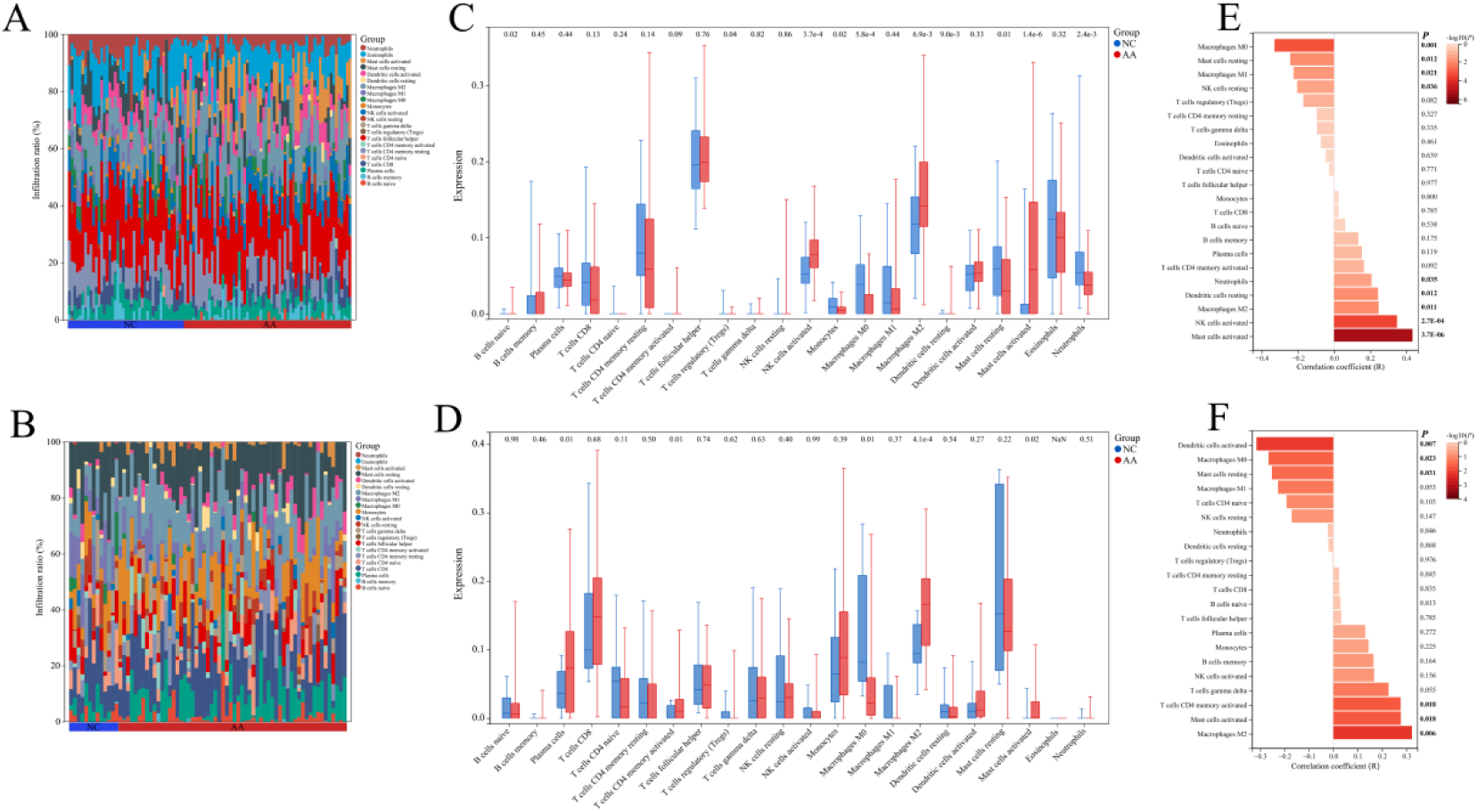
Immune infiltration analysis. (A, B) Barplot showing the infiltration levels of 22 immune cell types in each sample of the GSE67472 and GSE147878 datasets. (C, D) Violin plot showing the infiltration differences of immune cells between the AA and NC groups in the GSE67472 and GSE147878 datasets. (E, F) Bar plot showing the correlation between TFF3 expression and immune cell infiltration levels in the GSE67472 and GSE147878 datasets.

### 3.6 Inhibiting TFF blocks the activation of the PI3K/AKT signaling pathway and inflammation in asthma models

The Western blotting results show that the expression of p-PI3K and p-AKT in 16HBE cells induced by HDM was significantly increased (Supple Figure 9A,9B). RT-qPCR (Supple Figure 10A) with western blot (Supple Figure 10B) results showed that TFF3 expression level was significantly increased in HDM-induced 16HBE cells compared with the control group. The data indicated elevated TFF3 expression in the asthma model, which was consistent with our previous bioinformatics results. The data suggest that inhibiting TFF blocks the activation of the PI3K/AKT signaling pathway(Figure 8A). For further verification, we established an airway inflammation model of asthma in vitro by using HDM to induce 16HBEcells and HAM cells. RT-qPCR results showed a significant increase in the expression levels of IL-13, IL-25 and IL-33 in 16HBE cells induced by HDM compared to the control group (Figure 8B). The expression levels of IL-4, IL-5, IL-6 and IL-13 were significantly increased in HAM cells induced by HDM (Figure 8B). Data indicated that inhibiting TFF decrease the HDM-induced inflammatory index in the asthma model, resulting in airway inflammation.

**Figure 8.**
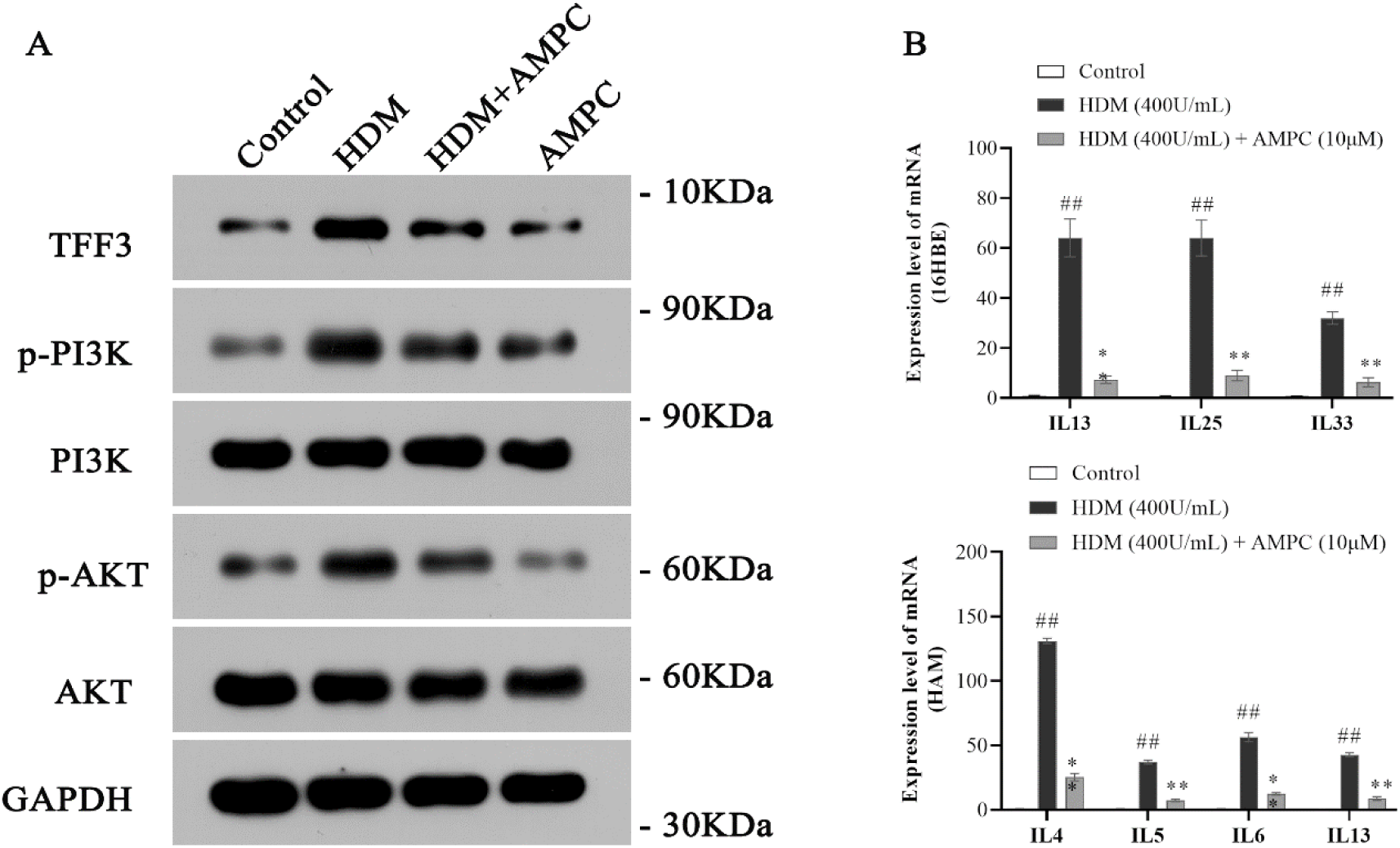
(A) Protein expression analysis in 400U/mL HDM-induced asthma model with or without 10μM AMPC (TFF3 inhibitor). TFF3, AKT, p-AKT, PI3K and p-PI3K expression in HDM-induced 16HBE cells were detected by western blotting. (B) Inflammatory factor expression analysis in asthma. Expression of IL-13, IL-25, and IL-33 mRNA in HDM-induced 16HBE cells werse detected by RT-qPCR. The expression of IL-4, IL-5, IL-6 and IL-13 mRNA in HDM-induced HAM cells was detected by RT-qPCR. ^##^*P*<0.01 vs Control group, \*\**P*<0.01 vs HDM group. Data are expressed as mean ± standard deviation, n=3.

## 4 Discussion

In this study, we adopted a variety of bioinformatics analysis methods to systematically investigate the expression pattern, regulatory network and function of TFF3 in asthma based on asthma-related transcriptome data, aiming to provide clues for in-depth research on the mechanisms of TFF3-mediated asthma pathogenesis. In addition, ROC curve analysis results showed that the AUC value of using TFF3 expression levels to diagnose asthma reached over 0.75, indicating its good diagnostic efficacy. Currently, the diagnosis of asthma mainly relies on clinical indicators such as medical history, signs, and lung function, lacking highly specific molecular markers [11]. Our results suggest that TFF3 may become a potential diagnostic biomarker for asthma.

Through differential analysis and WGCNA, we screened out 57 genes significantly associated with asthma phenotype and TFF3 expression. PPI network analysis further revealed that 53 genes had direct or indirect interactions with TFF3, suggesting that they may be major upstream or downstream factors regulated by TFF3 in the occurrence and development of asthma. We performed expression validation and molecular docking verification for TFF3 direct interaction genes. The results showed that 11 TFF3 direct interaction genes (AGR2, AKT1, ATP10B, CLDN10, CPA3, CXCL14, DEFB1, MS4A2, S100P, SCGB2A1, SPDEF) were significantly highly expressed in the airway epithelial tissues of asthma patients and were significantly positively correlated with TFF3 expression. Molecular docking analysis confirmed that TFF3 had good binding ability with the proteins encoded by these genes, providing structural biological evidence for their direct regulatory relationship. These results indicate that the 11 TFF3 direct interaction genes may be key regulatory factors for TFF3 involvement in the occurrence and development of asthma. Some genes have been reported to be related to the asthma process or TFF3 regulatory function. For example, AKT1 can induce bronchial smooth muscle cell proliferation, aggravating airway hyperresponsiveness and airway wall thickening [15], suggesting that TFF3 may be involved in airway smooth muscle remodeling by activating AKT1, but the specific regulatory effects need to be further investigated and confirmed.

we further verified the expression of TFF3 in asthma through in vitro experiments. We chose airway epithelial cells (16HBE cells) for the experiments to complete the construction of the cell model by using the HDM-induced airway inflammation model of asthma, and detected the inflammation-related indexes. Our study found elevated expression of TFF3 in an in vitro asthma model, which is consistent with results obtained from bioinformatics. There is evidence that TFF 3 can act on the epidermal growth factor receptor (EGFR) and then activate several downstream signaling pathways, including the PI3K/AKT pathway. TFF3 has been found to be involved in the activation of the PI3K/AKT signaling pathway, and TFF3 overexpression increased the level of phosphorylated AKT. It is suggested that TFF3 may affect the development of asthma by activating the PI3K/AKT signaling pathway, and further experiments are needed to demonstrate this.

Functional enrichment analysis results suggest that TFF3 may participate in the pathological processes of asthma by being involved in biological processes and signaling pathways such as cell migration, differentiation and proliferation, immune response, Fc epsilon RI signaling pathway, Sphingolipid signaling pathway, and Phospholipase D signaling pathway. Studies have shown that goblet cell differentiation and proliferation are one of the important pathological changes in asthma airway remodeling, leading to airway secretory function hyperactivity and mucus hypersecretion, aggravating airway obstruction [18]. TFF3 induces inflammatory responses and excessive mucus production in asthma by promoting goblet cell differentiation and proliferation [3]. The Fc epsilon RI signaling pathway plays a key role in IgE-mediated type I hypersensitivity reactions. Studies have shown that IgE levels are significantly elevated in asthma patients, and binding to Fc epsilon RI activates downstream signaling molecules such as Lyn and Syk, mediating mast cell hypertrophy and activation, releasing inflammatory mediators such as histamine and leukotrienes, causing bronchial smooth muscle contraction and airway hyperresponsiveness [19–21]. In addition, TFF3 can bind to IgG Fc binding protein (FCGBP) and activate the Fc epsilon RI signaling pathway, thereby inducing mucosal innate immune responses and promoting tracheal epithelial mucus secretion [22, 23]. The Sphingolipid signaling pathway is involved in regulating various biological processes such as cell proliferation, differentiation, apoptosis, and inflammation [24]. Studies have found that sphingolipid metabolic disorders are closely related to airway inflammation and airway remodeling in asthma [25]. Sphingolipid metabolites such as sphingosine and sphingosine-1-phosphate can promote airway smooth muscle cell proliferation and release of inflammatory factors such as IL-6 by activating signaling pathways such as MAPK [26, 27]. The Phospholipase D signaling pathway participates in processes such as cell proliferation, differentiation, migration, and inflammation by hydrolyzing phosphatidylcholine to produce phosphatidic acid and choline [28]. Studies have found that Phospholipase D activity is significantly elevated in lung tissues of asthmatic mouse models, promoting IL-13-induced airway hyperresponsiveness [29]. Therefore, we speculate that TFF3 may play an important role in airway inflammation and airway remodeling in asthma by regulating the above signaling pathways.

Immune cell infiltration is crucial in chronic airway inflammatory responses in asthma. The study found that in asthma patients’ airway tissues compared to healthy controls, the infiltration levels of Macrophages M2 and activated Mast cells were significantly increased, while that of Macrophages M0 was significantly decreased. Correlation analysis indicated that TFF3 expression levels were positively correlated with the infiltration of Macrophages M2 and activated Mast cells, and negatively correlated with the infiltration of Macrophages M0 and resting Mast cells. This suggests that TFF3 may be involved in airway inflammatory responses in asthma by regulating the infiltration of different subtypes of immune cells.

Macrophages are significant effector cells in asthma airway inflammation. They are categorized into M1 (classically activated) and M2 (alternatively activated) types based on activation status and function. Research indicates that M2 macrophages are markedly increased in airway tissues of asthma patients and have a crucial role in Th2-type airway inflammatory responses. [30]. Our study found that TFF3 was positively correlated with M2 macrophages and negatively correlated with M1 macrophages, suggesting that TFF3 may preferentially promote M2 macrophage polarization and recruitment, thereby aggravating Th2-type airway inflammatory responses. Mast cells are also key effector cells involved in the pathogenesis of asthma. Studies have shown that activated mast cells are significantly increased in bronchoalveolar lavage fluid and bronchial mucosal biopsy tissues of asthma patients [33–35]. Activated mast cells release various inflammatory mediators such as histamine, leukotrienes, and prostaglandins through cytoplasmic degranulation, causing bronchial smooth muscle contraction and increased vascular permeability, which is the initiating link of immediate hypersensitivity reactions in asthma [21]. In addition, mast cells can also secrete Th2 cell factors such as IL-4, and IL-13, as well as chemokines such as CCL5 and CXCL8, further aggravating hypersensitivity inflammation [36]. In our study, TFF3 was positively correlated with activated mast cells, indicating that TFF3 may have the effect of promoting mast cell activation.

This study confirms the close relationship between TFF3 and asthma pathogenesis. TFF3 may participate in asthma’s pathological processes by PI3K/AKT1 signaling pathways. It may also promote Th2-type immune responses by regulating macrophage polarization and mast cell activation. This provides important clues for understanding TFF3’s molecular mechanisms in asthma and new ideas for asthma diagnosis and treatment.

## 5 Conclusion

This study used bioinformatics methods and in vitro experiments to systematically explore TFF3 in asthma. Results showed TFF3 is highly expressed in asthma patients with good diagnostic efficacy. Multiple genes related to asthma and TFF3 expression were identified. PPI network analysis found potential interacting genes. These genes are involved in asthma-related processes and pathways. Molecular docking verified TFF3’s interaction with proteins. Immune infiltration analysis showed a correlation with M2 macrophages and mast cells. In vitro model verified TFF3’s high expression and its possible role through the PI3K/AKT signaling pathway.

In summary, this study used bioinformatics analysis and an in vitro asthma model to show TFF3’s important role in asthma pathogenesis across multiple levels. It deepens understanding of asthma’s molecular mechanisms and provides potential biomarkers and drug targets. Future studies are needed to further validate these findings and explore TFF3’s molecular mechanisms in asthma pathogenesis.

## Acknowledgements

None.

## Authors’ contributions

Fang, He collected data, performed experiments, analyzed data, and drafted the manuscript. Shu Wang performed cell experiments. Pengfei Huang performed analysis and interpretation for the data. Xiaoyun Fan designed and supervised the study, and modified the manuscript. The final manuscript for publication is read and approved by all authors.

## Data availability

The datasets used in this study are available from GEO database (https://www.ncbi.nlm.nih.gov/geo/) at NCBI, STRING database (https://cn.string-db.org/), the DAVID database (https://david.ncifcrf.gov), the CIBERSORTx database (https://cibersortx.stanford.edu/). The HDOCK protein-protein docking server (http://hdock.phys.hust.edu.cn/) was used for molecular docking analysis.

## Declarations

### Ethical Approval

This study did not involve any human participants or animal experiments. Therefore, ethical approval was not required.

### Consent for publication

Not applicable.

### Clinical trial number

Not applicable.

